# Interpreting the fossil record and the origination of birds

**DOI:** 10.1101/2022.05.19.492716

**Authors:** Nicholas M. A. Crouch

## Abstract

The fossil record is essential for understanding when lineages originate and their pace of diversification. However, numerous taphonomic biases in the fossil record can hinder interpretation, creating discord between palaeontological and phylogenetic estimates of clade origination dates. Here, I use the recently published Bayesian Brownian Bridge method to infer the age of birds using occurrence data from the Paleobiology Database. I also estimate the age of the speciose sub-clade Telluraves to compare age estimates with previous tip-dating analyses of the same group. Analyses of all birds show place the root age approximately 100 Ma, approximately 26 Myr before the oldest fossil occurrences. Increasing the time bin size from 2.5 to 5 Myr produced significantly older and less precise estimates. Divergence estimates for Telluraves were strikingly consistent with tip-dating analyses, placing origination of the group in the latest Cretaceous. Although these dates are consistent with a hypothesis of Mesozoic origination and Cenozoic diversification, significant diversification was estimated before the end-Cretaceous mass extinction suggesting analyses using pooled species counts may produce spurious results. Overall, these analyses provide further evidence to a growing consensus that several major avian lineages survived the end-Cretaceous mass extinction before diversifying into the most speciose extant tetrapod radiation.

---

The fossil record provides invaluable insight into macroevolutionary trends, yet these inferences are conditional on accurately dating the divergences between lineages. Dating lineages can be achieved in two principal ways: by strict interpretations of the fossil record where by the first appearance of a taxonomic group defines its age, and by phylogenetic methods where by the molecular clock is calibrated by placing priors on node ages or by the including extinct taxa directly in the reconstruction as terminal taxa. Although these two approaches can sometimes produce harmonious results, the fossil record is well documented by having varying taphonomic biases which frequently hinder interpretation. Preservation biases can be induced through a variety of processes. For instance, the fossil record may show temporal variation in diversity due to variation the number of fossil bearing formations, and the volume of rock encompassed by these formations [1—6]. The preservation potential of the rock record can also vary according to different stages of continent formation [7]. Preservation biases may also be lineage-specific. For example, aspects of species’ ecology may result in a greater probability of being fossilized, such as large body size, and species found in different habitats may also vary in their preservation rates [8]. Therefore, reconciling differences between ‘strict’ interpretations of the fossil record and other records must necessarily take these variations into account.

While angiosperms have the notoriety of being Darwin’s “abominable mystery” [9], dating the origination and major divergences within birds has also been a long-standing debate. Broadly, molecular studies have historically suggested a Cretaceous origin, preceding the K-Pg boundary [10—19]. However, such an early origin is highly controversial [20] given the general absence of specimens from the early Cenozoic [21, 22]. A possible explanation for the discrepancy is an over-estimation of rates of molecular evolution early in the history of the group due to strong selection against larger-bodied taxa at the end-Cretaceous mass extinction [23]. A recent tip-dating study, where the fossil record is included directly into the phylogenetic reconstruction along with molecular data [24], placed the origination of a large clade of birds (Telluraves) much later in the Cretaceous (95% HPD: 69.9–82.8 Mya) with diversification following the end-Cretaceous mass extinction [25]. While this study allowed for comparatively greater reconciliation between the two approaches, this analysis still resulted in significant gap between the crown age and the oldest fossils of the group, leaving open the question of when these enigmatic taxa first appeared.

In this study, I analyze the fossil record of birds (crown Aves) using the recently published Bayesian Brownian bridge (BBB) method [26]. This method uses information on the fossil record and extant diversity to estimate time of origination of the clade, its diversification history, and sampling rate. Moreover, it specifically accommodates for clades with poor fossil records, such as birds. I use almost ten thousand occurrences for birds entered into the Paleobiology Database in conjunction with revised estimates of the numbers of extant species. Additionally, I estimate the age of a sub-clade of birds – Telluraves, a large clade of predominantly arboreal species which covers approximately two-thirds of extant diversity – to contrast BBB estimates with those derived from a previous tip-dating study [25].

## Materials and Methods

I obtained data on the avian fossil record from the Paleobiology Database [27]. Although the database contains imperfections, it still represents an enormous resource on fossil occurrences of clades across the Tree of Life. Additionally, I performed these analyses at wide taxonomic scales (i.e. all birds and the clade Telluraves) which should accommodate for some of the more well-known idiosyncrasies of the avian fossil record, such as the tendency for new specimens to be assigned to new genera and species. I downloaded occurrence data using the R package paleobioDB [28] on January 29th, 2021. I subset the data to only contain species-level records, and also excluded enantiornithine (e.g. Alexornithiformes) and non-avian theropod orders (e.g. Omnivoropterygiformes) from all analyses. All taxa removed from the analysis are documented in the supplementary material. I converted the occurrence data into BBB format using custom R scripts to construct family-level occurrence data (following Silvestro et al. [26]) which I then summed to create data for higher taxonomic units. Although this pooling implies equal preservation potential [26], accurate dating of birds almost certainly requires consideration of extinct families present in the late Cretaceous and early Cenozoic which cannot yet be accommodated in BBB. In the supplement I provide a custom R function to allow easy replication of these analyses for any taxonomic group, with this function performing the following steps. First, it excludes of any specimens that cannot be identified to family level. Next, it makes a complete list of all families using data on extant taxa plus those which only appear in the fossil record (for example the Zygodactylidae in birds). Finally, within specified time bin intervals, counts of each family is made across the recorded duration of the clade.

I analyzed three taxonomic subsets of birds: all crown species plus stem taxa (e.g. *Hesperornis*), all crown species excluding stem taxa, and finally orders from the clade Telluraves to compare the results with previous tip-dating efforts [25]. This previous analysis by Crouch et al. [25] did not sample every order within Telluraves; therefore, I only analyzed the same eight orders to facilitate the most accurate comparison. Within these three taxonomic subsets I created occurrence data in 2.5 and 5 Myr time bins respectively, giving a total of six BBB runs. I left the prior on the root age as a wide uniform distribution with a maximum bounds at 300 Ma so as to not bias the results. The most widely used estimate of extant avian diversity is that of 9,993 species from Jetz et al. [17]. However, ongoing taxonomic revision, particularly molecular analyses, suggest this to be an under-estimate [29]. Therefore, I ran the analyses specifying the most conservative estimate from Barrowclough et al. [29] of 15,845 species. For the analysis of Telluraves orders I expanded the taxonomic sampling of [17] by the same amount, resulting in a taxonomic diversity of 11,100 species. I ran all analyses for 500,000 generations, sampling every 200, and specifying a non-constant sampling rate through time. I estimated whether the analyses had reached convergence by visualizing the posterior distribution of the parameter estimates from each model. I discarded 50% of the posterior sample before visualizing the distribution of parameter estimates.

## Results

I downloaded 14,668 occurrences from the Paleobiology Database, representing 141 out of the 194 families (73%) present in the taxonomy of [17]. There were an additional 124 families present in the Paleobiology Database taxonomy not present in the taxonomy of extant species, resulting in 318 unique families analyzed. Removing species outside of Aves resulted in 9891 occurrences to be analyzed. The distribution of fossil ages is extremely skewed, with 90% of the records being confined to within the last 10 Myr. The five families with the greatest numbers of occurrences in the data are either associated with aquatic habitats (Procellariidae, Anatidae, Rallidae), larger body size (Accipitridae), and/or high extant abundance (Columbidae).

Estimated root ages for the analysis of all birds was significantly affected by time bin size (Figure 1). When 2.5 Myr bins were used the root age was estimated in the mid-Cretaceous, with significant overlap in the analyses including an excluding stem taxa. Analyzing the data using 5 Myr bins increased the overall root age, as well as discrepancy between analyses including and excluding stem taxa, pushing the root age back into the Jurassic. This increased the gap between the oldest fossil record and the estimated root age from 26 to 113 Myr for the analysis excluding stem taxa, and from 34 to 84 Myr for the analysis including stem taxa. Differences in root ages were driven by changes in the *a* parameter, which describes the exponential rate at which the sampling rate changes (Figure 1). The estimated root age for Telluraves was strikingly similar to those recovered by the tip-dating analysis of Crouch et al. [25], but with greater uncertainty (HPD 55 – 92 Ma). This analysis resulted in a gap of 21 Myr between the oldest specimens and the root age. Changing the size of the time bins from 2.5 to 5 Myr intervals also resulted in less precise estimates for the estimated root age (Figure 1). All analyses recovered the vast majority of avian diversification in the Mesozoic (Figure 2), consistent to when the analyses of Telluraves were analyzed separately.

**Figure 1:**
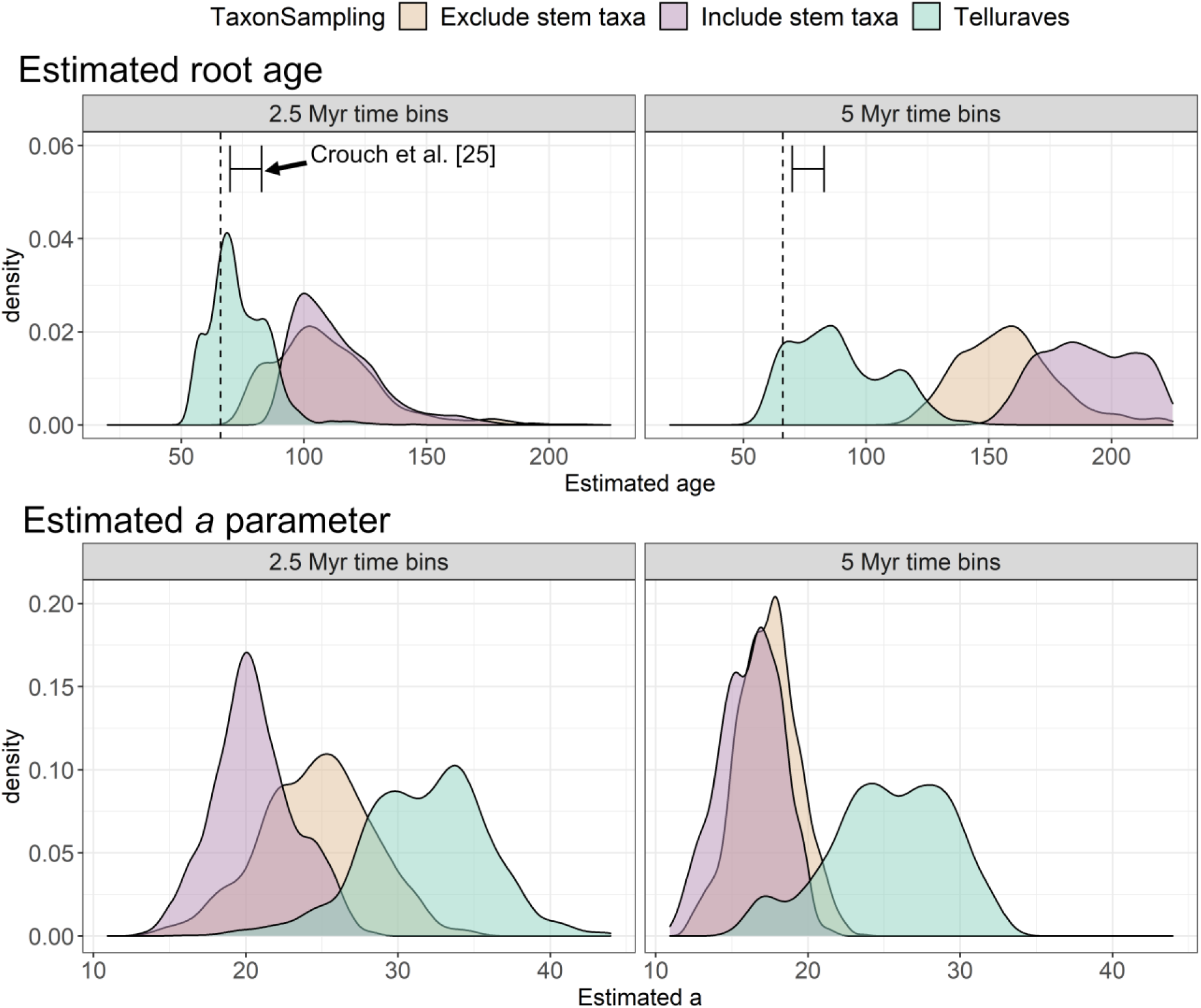
Comparison of select model parameters estimated using Bayesian Brownian bridge [26]. The top row shows the estimated root age, and bottom row shows the *a* parameter, describing the rate of exponential increase in the sampling rate as a function of time. In all panels colors indicate different taxon sampling regimes. The panels correspond to the size of the time bins used in the respective analyses. The error bars shown in the two panels show the 95% highest probability density of the age of Telluraves estimated using tip-dating [25].

**Figure 2:**
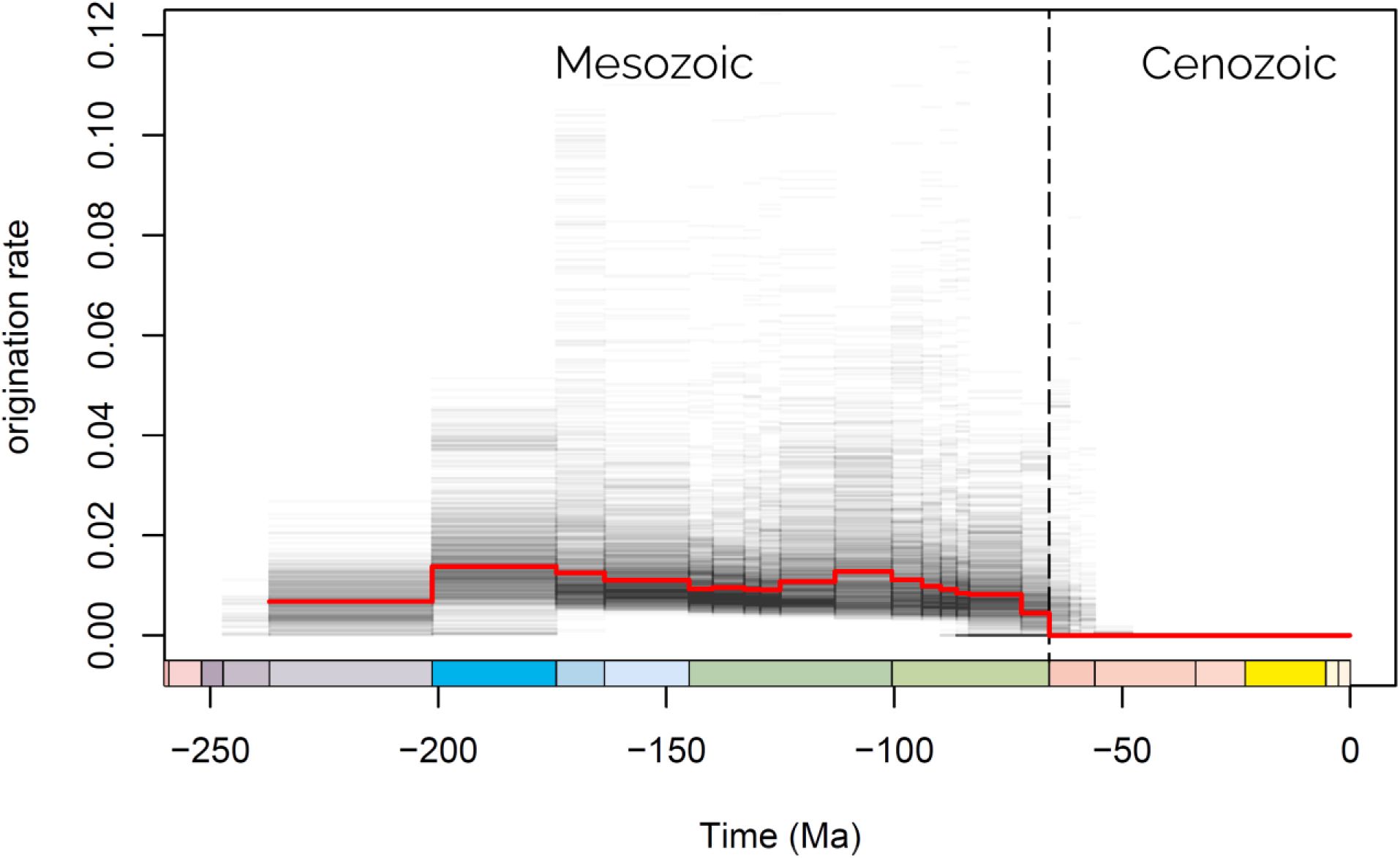
Bayesian Brownian Bridge estimates of avian diversification.

## Discussion

Accurate inference of evolutionary processes is conditional on accurate dating of when lineages originate. Here I use the recently published BBB method [26] to estimate the age of birds, as well as the sub-clade Telluraves to compare to tip-dating estimates [25]. These results show first that the time bin size for BBB analyses should likely not exceed 2.5 Myr. Although the method has been shown to be robust against even extremely poorly sampled fossil records [26], increasing the size of the time bins in this manner reduces the precision of root age estimates, as well as producing unrealistically old root age estimates for birds (Figure 1). The increase in estimated age is influenced by changes to the *a* parameter between the three analyses. This parameter was significantly shallower in the analyses of all birds, indicating a less pronounced slope to the exponential distribution. A shallower curve may be required to explain the extremely skewed distribution of fossil ages, which is less pronounced if the analysis is concentrated towards the present as in the analysis of Telluraves. A contribution of the distribution of avian fossil ages and larger time bins may therefore be causing the estimated age to be stretched back to the Jurassic. Therefore, as well as recommending a maximum time bin age of 2.5 Myr, these results suggest that current implementation of BBB should be restricted to the analysis of groups which have fossil occurrences spanning the majority of the estimated age of the group, rather than pooling extinct groups into a single analysis.

The BBB estimate of the age of Telluraves was remarkably consistent with the tip-dating analysis of Crouch et al. [25]. This previous analysis aimed to capture as much taxonomic variation in the fossil record as possible within the clade, with this resulting in a notably sparse morphological matrix (see [25] for full details). Although this sampling rate was not unusual for a study of its size, it raised concerns that it may have affected both estimated topologies and rates of evolution, potentially distorting the overall age of the group. These analyses show that, even where the amount of morphological data available for extinct species is limited, tip-dating is a powerful method for estimating the divergences between lineages. The comparatively greater uncertainty in the age estimates from BBB could be due to the comparatively reduced amount of data available for dating the reconstructions (i.e. genetic and morphological data as well as the ages of specimens). Nevertheless, the BBB method will likely be an invaluable tool for regions of the Tree of Life where construction of morphological matrices is either not possible or significantly challenging.

BBB estimates of avian diversification suggest this process was significantly faster in the Mesozoic than Cenozoic (Figure 2). A similar pattern is also recovered when from the analyses of only Telluraves. This result is concerning, given that it is highly unlikely crown birds both originated and diversified in the Mesozoic; instead, the ecological release following the end-Cretaceous mass extinction is considered a more plausible process [30, and references therein]. This result may be driven by two factors: the paucity of the record in deep time or the pooling of extinct families into a single analysis. Given BBB was designed to handle highly incomplete data, the latter seems the more likely explanation. Therefore, these results suggest that continued modification is required to handle extinct lineages, which will only serve to make BBB a more powerful tool for inferring historical processes.

Resolving the age of birds has been a long-standing debate. However, aided by the continued discovery of specimens, especially from the Cretaceous period, there is an ‘emerging consensus’ to form regarding the evolution of the group [30]. For analyses within the crown, these results fit within this theory – that a limited number of avian lineages were able to survive the end-Cretaceous mass extinction, possibly due to factors such as their incubation mode [31], followed by explosive diversification in the Cenozoic likely aided by the removal of competing taxa. This work serves to strengthen this hypothesis by demonstrating that palaeontological estimates of avian diversification can be congruent with phylogenetic analyses.

## Acknowledgments

I would like to thank D. Silvestro for comments on running BBB, and K. Collins for comments on the manuscript.

## Data availability

The data for this study were downloaded from the Paleobiology Database. Code used in the analyses of these data and BBB output files are provided on Dryad (https://doi.org/10.5061/dryad.j0zpc86cx).

